# Modelling the effects of PPARβδ of innate inflammatory responses in lung tissues

**DOI:** 10.1101/2020.06.07.138867

**Authors:** Noelia Perez Diaz, Lisa A Lione, Victoria Hutter, Louise S. Mackenzie

**Affiliations:** School of Life and Medical Sciences, University of Hertfordshire, Hatfield, AL10 9AB, UK; School of Pharmacy and Biomolecular Sciences, University of Brighton, Brighton, BN2 4GJ

**Keywords:** Lung, nuclear receptor, gene transcription, inflammation, molecular docking, PPARβ/δ, molecular switch, induction, trans-repression, inflammation, lung, pulmonary artery, GW0742, GSK3787, docking

## Abstract

Peroxisome proliferator activated receptor beta/delta (PPARβ/δ) is a nuclear receptor ubiquitously expressed in cells whose signaling controls inflammation and metabolism. However, there are great discrepancies in understanding the role of PPARβ/δ, having both anti- and pro-effects on inflammation. Understanding the PPARβ/δ mechanism of action may provide new molecular mechanisms for treating a variety of inflammatory-related diseases.

We studied the PPARβ/δ-regulation of LPS-induced inflammation of pulmonary artery, bronchi and parenchyma from rat, using different combinations of agonists (GW0742 or L-165402) and antagonists (GSK3787 or GSK0660). LPS-induced inflammation is largely regulated by PPARβ/δ in the pulmonary artery, but it is a minor factor in bronchi or parenchyma. Agonists do not significantly inhibit inflammation, but activates the PPARβ/δ induction mode of action. Surprisingly, co-incubation of the tissue with agonist plus antagonist shows anti-inflammatory effects and switches the PPARβ/δ mode of action from induction to trans-repression, indicating that the PPARβ/δ induction mode of action is pro-inflammatory and the trans-repression anti-inflammatory. Us of Computational chemistry methods indicates that PPARβ/δ agonists are predicted to form polar interactions with the residues His287, His413 and Tyr437 whilst PPARβ/δ antagonists form polar interactions with the residues Thr252 and Asn307. Further, our modelling indicates favorable binding energies and the feasibility of simultaneous binding of two ligands in the PPARβ/δ binding pocket. In summary, this study provides novel insight into the complex relationship between ligand binding profiles and functional outcomes in a rat lung inflammation model, which will help inform the design of novel therapies for inflammatory lung diseases.

## Introduction

PPARβ/δ are ligand dependent transcription factors that belong to the nuclear receptor family (1). They are ubiquitously expressed in all cells tested (2) and control key biological functions such as lipid metabolism, glucose metabolism, immune response, inflammation, cell proliferation, cell migration, apoptosis and carcinogenesis (3-5). Consequently, PPARβ/δ is emerging as a therapeutic target for the treatment of disorders associated with metabolic syndrome, although there are no marketed drugs specifically targeting PPARβ/δ yet.

Research reported mainly on animal models such as mice, rats or rhesus monkeys indicates that PPARβ/δ agonists result in a number of favorable pharmacological effects including reduced weight gain, increased metabolism in the skeletal muscle and cardiovascular function, suppression of atherogenic inflammation and improvement of the blood lipid profile, which are common abnormalities in patients with metabolic syndrome (3,6,7).

These encouraging results led to the first clinical trials on humans. Glaxo Smith Kline (GSK) developed the agonist GW501516 (Endurobol), a promising compound that completed proof-of-concept clinical trials successfully for dyslipidemia (8) and hypocholesteremia (9). However, further studies showed a suspected link with tumor development (10,11), and any further clinical trial with GW501516 was suspended.

Nevertheless, the interest in PPARβ/δ continues and in the last few years several compounds targeting PPARβ/δ were developed, although just a few of these molecules reached clinical trials. One of them was MBX-8025, which reached Phase II clinical trial for patients with non-alcoholic steatohepatitis and primary sclerosing cholangitis; unfortunately, this study was terminated early when patients developed early signs of liver damage as indicated by liver biopsy (12).

PPARβ/δ can be activated by numerous endogenous ligands such as eicosanoids, fatty acid, metabolites derived from arachidonic acid and linoleic acid (13-15) as well as exogenous synthetic ligands like GW0742, L-165041, MBX-8025 and GW501516, and can also be repressed by the two synthetic antagonists GSK3787 and GSK0660.

After ligand activation, PPARβ/δ regulates genes by two different mechanisms, induction and trans-repression. In the induction mode, PPARβ/δ forms a complex with the retinoid X receptor (RXR) and together, as a heterodimer, binds the promoter of the target genes (PPRE). In the absence of ligand, co-repressor proteins and histone deacetylases (HDACs) are bound to the heterodimer which tight the chromatin and prevent it from binding to the PPRE sites (16). The presence of ligand induces a conformational change of PPARβ/δ which promotes the binding of co-activators, releases the co-repressor proteins, induces histone acetylation and methylation and finally allows the transcription of the target genes (17,18).

In the trans-repression mode PPARβ/δ regulates gene expression in a PPRE-independent manner through the regulation (mostly suppression) of other transcription factors, including nuclear factor-κB (NF-κB) (19), activator protein (AP-1) (20) and B cell lymphoma 6 (Bcl6) (21). However, there are great discrepancies in the literature about the effect of the ligand-activation of PPARβ/δ in the cell, and both pro- and anti-effects in inflammation (22,23), cell proliferation (24,25) and migration (26,27) have been reported. PPARβ/δ receptor appears to be a sensitive molecular switch that has both endogenous and exogenous ligands which control cellular function through changes in very small concentration range. Added to this, in any cell or tissue, the activity of PPARβ/δ may also depend on its promoter activity and relative expression, as well as presence and activity of co-repressor and co-activator proteins. It has been shown that GW0742 is capable of behaving as an agonist activating the transcription pathway at lower concentrations (nM) and antagonist inhibiting this effect at higher concentrations (μM) (28). In the same line, a study in a model of systemic inflammation in mice showed that higher doses of GW0742 (0.3 mg/kg) triggered a pro-inflammatory response, whereas a lower concentration (0.03 mg/kg) showed an anti-inflammatory trend, although without a significant difference (29).

Understanding how PPARβ/δ switches between induction and trans-repression mode of action and how this determines cellular function is of great interest and may provide new molecular targets for treating a variety of inflammation-dependent diseases, including atherosclerosis, diabetes, and cancer. Whether differences in endogenous and exogenous ligands induce slightly different genes in different cells is a great possibility, and one that has not been so far explored. The question that needs to be addressed is not whether activation leads to proliferation and cancer, but whether the type of agonist activation and the subsequent molecular control can be adjusted to place the cell into a non-proliferative and non-inflammatory state of gene expression. A parallel argument has been made on the cause of the side effects of glucocorticoids; the type of ligand determines whether the glucocorticoid receptor forms a homodimer, and the subsequent type of trans-repression and transcriptional control exerted leads to change in cellular function (30). Therefore, we hypothesized that PPARβ/δ acts as a molecular switch between induction and trans-repression and depending on which mechanism is triggered, it will have pro- or anti-effects.

In order to address these fundamental questions, we investigated the varying effects of PPARβ/δ ligands on inflammation in three isolated tissues within the lung: pulmonary artery, bronchi and parenchyma. Using marker genes to indicate the level of induction and trans-repression we showed how different tissues respond, and using computational docking of the PPARβ/δ with various combinations of the ligands, have insight that might explain the varying effects on the resulting changes in gene transcription that leads to the range of cellular functional changes.

## Results

### PPARβ/δ regulation of LPS-induced lung tissue inflammation

Three main tissues were identified in the lung; pulmonary artery, bronchi and parenchyma, and the expression of PAPRβ/δ was verified in all three tissues (**Supplementary Figure 1**). NO production from pulmonary artery, bronchi and lung parenchyma was measured after 8 h, 20 h and 24 h of incubation (**Figure 1**), and indicates that the effects of the treatments for all the tissues tested are greatest at 24 h of incubation.

**Figure 1.**
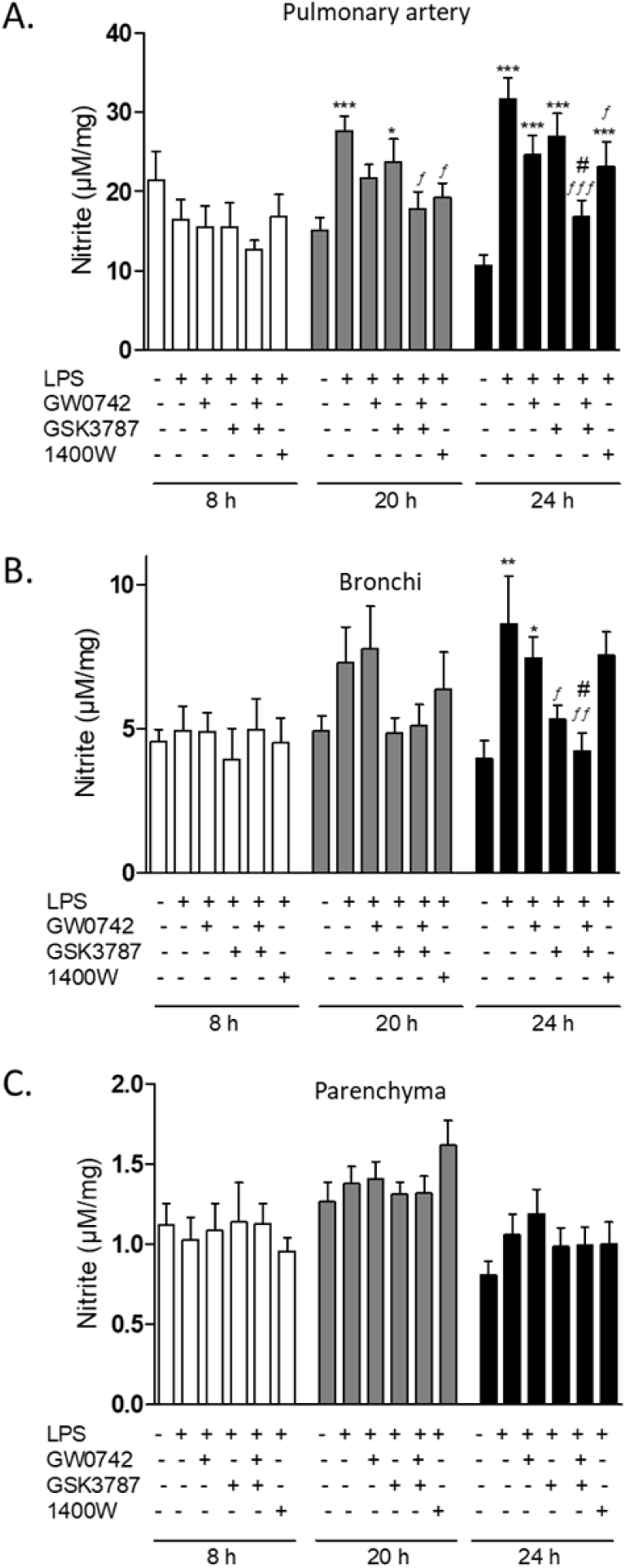
NO production by lung tissues over the time. A) Pulmonary artery (n=13), B) bronchi (n=9) and parenchyma (n=14) were dissected, cut into six pieces and incubated under six different treatments of different combinations of 1 µg/mL LPS, 100 nM GW0742, 1 µM GSK3787, 10 µM 1400W; the NO production was measured at 8 h, 20 h and 24 h of incubation. Significant difference between treatments was analyzed by two-way ANOVA followed by Bonferroni post-hoc test and the data are presented as mean ± standard error of the mean. *=P<0.05, **=P<0.01, ***=P<0.001 compared with vehicle; *f*=P<0.05, *ff*=P<0.01, *fff*=P<0.001 compared with LPS; #=P<0.05 compared with LPS+GW0742.

The pulmonary artery rings used for the time-line incubations were too small to use for further experiments, therefore the same experiment was repeated using bigger rings and NO and IL-6 production was measured after 24 h incubation (**Figure 2**). LPS significantly increased NO by 4 to 10-fold (**Figure 2A, 2B**), as expected. In the presence of PPARβ/δ ligands GW0742 and GSK3787 (**Figure 2A**) or L-165041 and GSK0660 (**Figure 2B**), nitrite production is not significantly different than LPS alone. In contrast, the combination of agonist GW0742 and antagonist GSK3787, or L-165041 and GSK0660 (**Figure 2A, 2B**) significantly reduces NO production. This indicates that the co-incubation with the agonist-antagonist attenuates the LPS-induced NO production more effectively. To confirm the inflammation IL-6 was measured by ELISA (**Figure 2C, 2D**). Again, LPS increases IL-6 production by 10-fold, and the PPARβ/δ ligands GW0742, L-165041 and GSK0660 do not have a significant effect on their own. However, the antagonist GSK3787 significantly decreases IL-6 production by half, as well as the co-incubation with GW0742-GSK3787 or L-165041-GSK0660.

**Figure 2.**
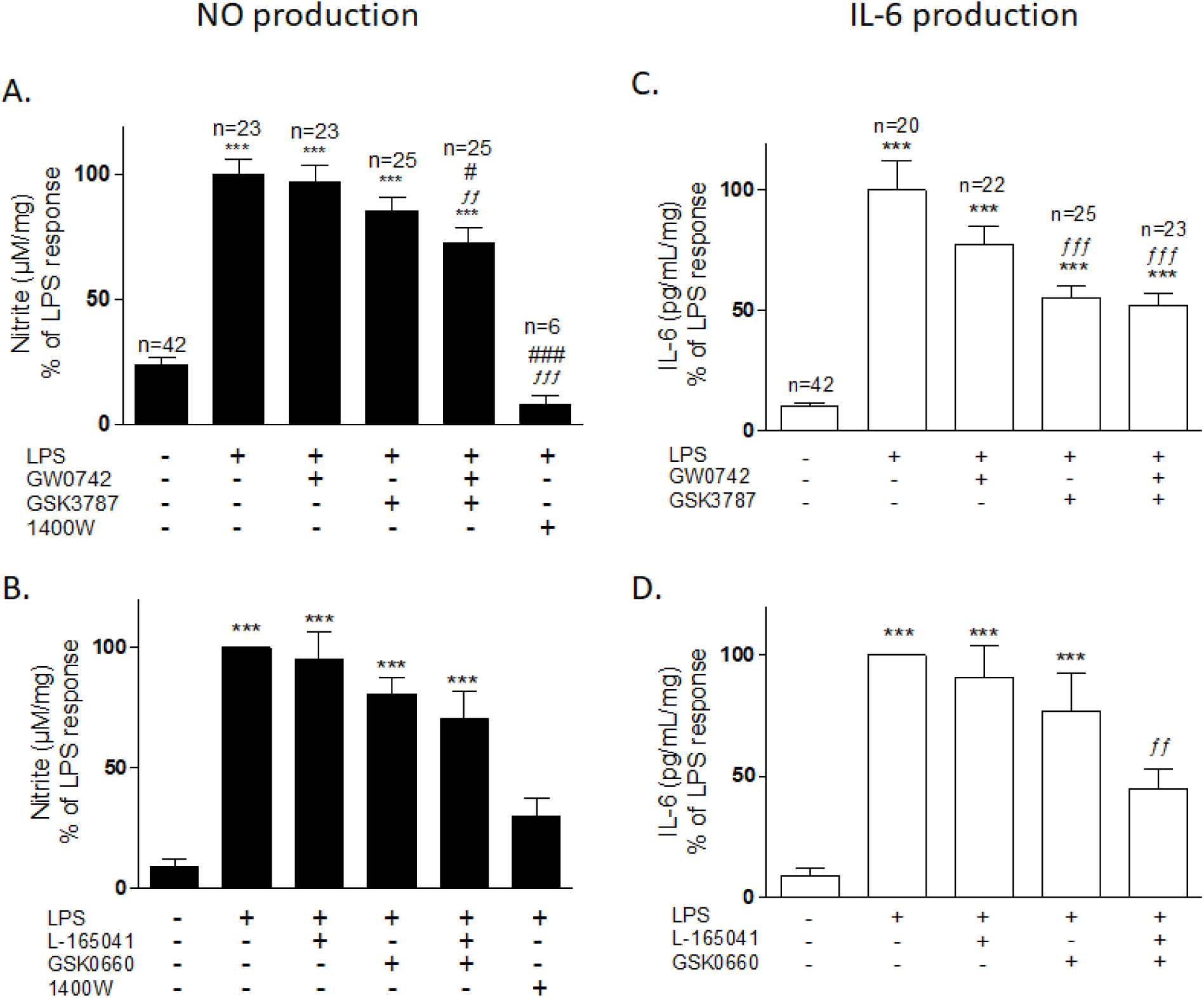
NO and IL-6 production by pulmonary artery. Rat pulmonary artery rings were treated with two combinations of PPARβ/δ agonist-antagonist and iNOS inhibitor 1400W: A-C) 100 nM GW0742 – 1 µM GSK3787 – 100 µM 1400W B-D) 1 µM L-165041 – 1 µM GSK0660 – 100 µM 1400W. NO and IL-6 production was measured after 24 h and normalized with LPS: A) The average of NO production in LPS was used for the normalization of the data and the number of samples per treatment is written at the top of the bar; B) each experiment was normalized with its own LPS treatment (n=9). C) The average of IL-6 production with LPS treatment was used for the normalization of the data and the number of samples per treatment is written at the top of the bar; D) each experiment was normalized with its own LPS treatment (n=9). Significant difference between treatments was analyzed by one-way ANOVA followed by Bonferroni post-hoc test and the data are presented as mean ± standard error of the mean. ***=P<0.001 compared with vehicle; *ff*=P<0.01, *fff*=P<0.001 compared with LPS; #=P<0.05, ###=P<0.001 compared with LPS+GW0742.

The same experiment was performed in bronchi with similar profile to pulmonary artery (**Supplementary Figure 2**); LPS increases the production of NO from 2- to 4-fold with LPS, which is decreased in the presence of the co-incubation with PPARβ/δ agonist and antagonist, although this reduction showed not to be statistically significant. LPS also increases IL-6 production 2 to 4-fold, but this increment is not significantly affected by the PPARβ/δ ligands used.

In parenchyma, LPS increases NO and IL-6 production by 2-fold and 3-4-fold respectively (**Supplementary Figure 3**). The PPARβ/δ ligands do not have any significant effect on LPS-induced NO, although the same pattern of NO reduction when agonist and antagonist are present at the same time is followed. Similarly, the PPARβ/δ ligands on their own do not have an effect on LPS-induced IL-6 production, however, the co-incubation with GW0742 and GSK3787 significantly reduces IL-6 by 1-fold.

### PPARβ/δ molecular switch in pulmonary artery

To understand the PPARβ/δ molecular switch, marker genes for the induction and trans-repression mode of action were chosen, and the transcription/repression of these marker genes were linked to the pro- and anti-inflammatory responses of the tissues tested. With that aim, *Pdk-4* and *Angptl-4* were chosen as markers of the induction mode of action and *Timp-1, Serpine-1* and *Id2* were chosen as markers for the trans-repression mode of action. The performance of the primers used were tested in the tissues and the data is summarized in **Supplementary Table 1**.

In pulmonary artery, GW0742 significantly increases the transcription of *Pdk-4* by 7-fold (**Figure 3A**) and *Angptl-4* by 3-fold (**Figure 3B**), which is inhibited by the antagonist GSK3787. The transcription of *Timp-1* (**Figure 3C**) and *Serpine-1* (**Figure 3D**) is increased by LPS in the pulmonary artery, and the agonist-activation of PPARβ/δ blocks the transcription by 3-fold compared to LPS. Interestingly, the treatment with LPS+GSK3787 increases the transcription of *Timp-1* which is again repressed by the treatment with LPS+GW0742+GSK3787. LPS does not induce the expression of *Id2* in the pulmonary artery, and similarly its expression is not affected by the PPARβ/δ ligands (**Figure 3E**).

**Figure 3.**
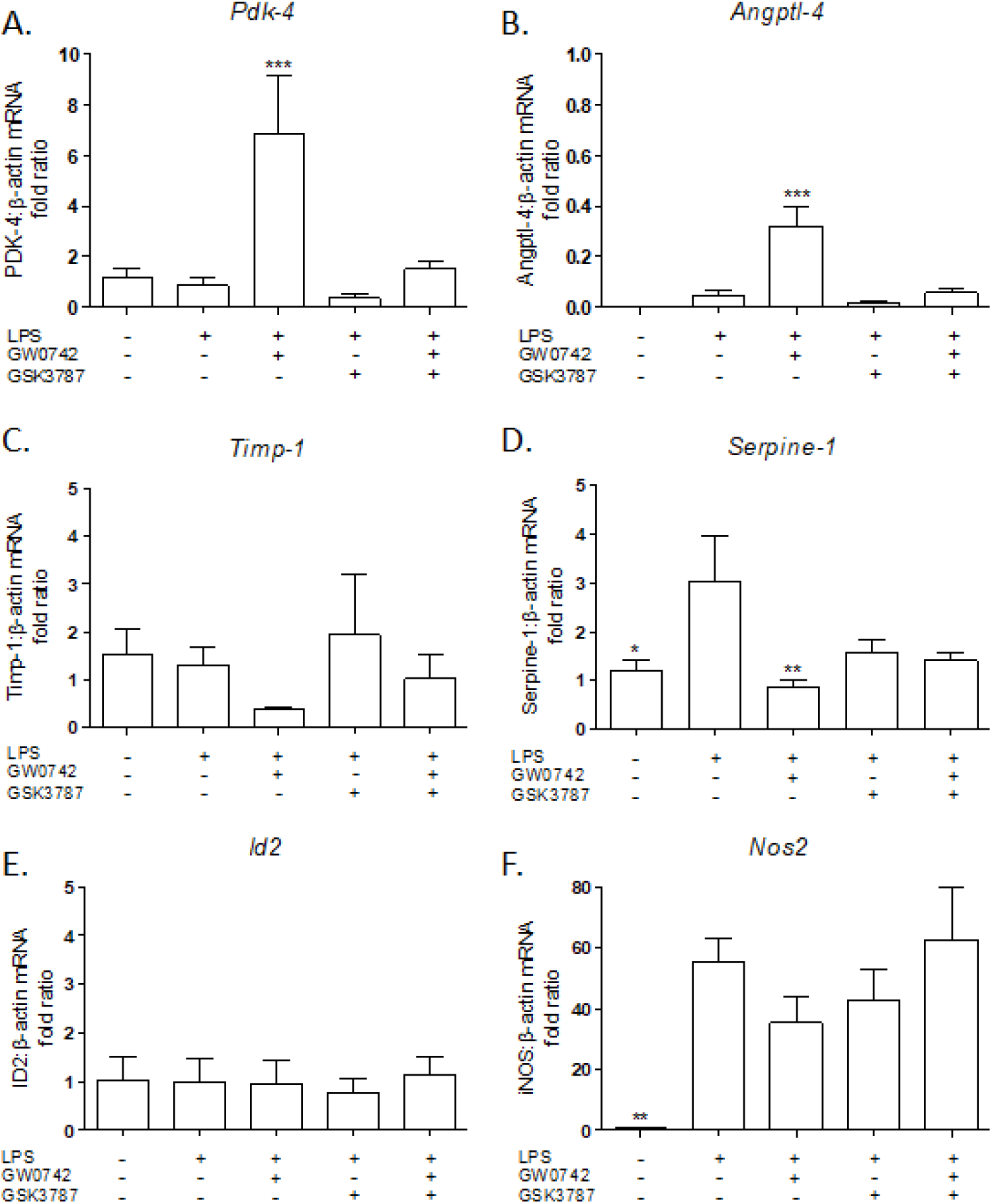
Expression of genes regulated by PPARβ/δ in the pulmonary artery. The expression of different PPARβ/δ target genes was measured after 24 h incubation under five different treatments: vehicle (0.01% DMSO); 1 µg/mL LPS; 1 µg/mL LPS + 100 nM GW0742; 1 µg/mL LPS + 1 µM GSK3787; and 1 µg/mL LPS + 100 nM GW0742 + 1 µM GSK3787 (n=4-5). Relative quantitation was calculated with the comparative CtΔΔ method and normalized against β-actin as an endogenous control. Significant difference between treatments was analyzed by one-way ANOVA followed by Dunnett’s post-hoc test and the data are presented as mean ± standard error of the mean. ***=P<0.001, **=P<0.01 and *=P<0.05 compared with LPS.

The expression of *Nos2*, the gene that encodes for iNOS, is significantly increased by LPS, and its expression is inhibited by GW0742 (**Figure 3F**), although this inhibition is not significant. Interestingly the treatment with LPS+GW0742+GSK3787 does not affect the expression of *Nos2*. Similar to pulmonary artery samples, GW0742 increased the expression of *Pdk-4* by 2-fold, although this increment is not significant, and the presence of GSK3787 inhibited its expression (**Supplementary Figure 4A**).

*Angptl-4* did not amplify in bronchi samples and the expression of *Timp-1, Serpine-1* and *Id2* showed not to be regulated by the treatments (**Supplementary Figure 4**). The transcription of Nos2 is significantly increased by LPS and to more extent by GW0742. The presence of GSK3787 significantly reduces the transcription of *Nos2* compared to LPS+GW0742, although it is still significantly higher than control (**Supplementary figure 4E**).

GW0742 increases the transcription of *Angptl-4* by 3-fold in lung parenchyma, however the rest of the marker genes tested were not affected by the PPARβ/δ ligands (**Supplementary Figure 5**).

### Computational Chemistry: PPARβ/δ Docking analysis

#### Docking of one PPARβ/δ ligand

The PPARβ/δ-LBD crystal structure 3TKM has an X-ray resolution of 1.95 Å and was co-crystallized with GW0742, the same agonist that was used during the development of this project, therefore this structure was chosen for our docking experiments.

The two PPARβ/δ agonists used in the previous experiments GW0742 and L-165041 as well as the two antagonists GSK3787 and GSK0660 were docked into the crystal structure of the LBD of PPARβ/δ. The best eight hits were analyzed by Pymol to identify the residues that form polar interactions with each of the different poses of the ligands (**Table 2**).

**Table 2.** Best eight docking hits of four ligands into PPARβ/δ (PBD: 3TKM). In green are the residues more likely to bind agonists and in red the residues more likely to bind antagonists

| Best fit | Agonists |  |  |  | Antagonists |  |  |  |
| --- | --- | --- | --- | --- | --- | --- | --- | --- |
|  | GW0742 |  | L-165041 |  | GSK3787 |  | GSK0660 |  |
|  | Affinity (Kcal/mol) | Aa with polar interactions | Affinity (Kcal/mol) | Aa with polar interactions | Affinity (Kcal/mol) | Aa with polar interactions | Affinity (Kcal/mol) | Aa with polar interactions |
| 1 | -11.1 | His287<br>His413<br>Tyr437 | -8.7 | His287<br>His413<br>Tyr437 | -9.1 | Thr252<br>Asn307 | -8.6 | Arg248<br>Thr252<br>Ala306<br>Asn307 |
| 2 | -10.8 | Thr253<br>His287<br>His413<br>Tyr437 | -8 | Met192<br>Thr252<br>Thr256<br>Ile290<br>Ala306 | -8.9 | Thr252 | -8.3 | Arg248<br>Thr252<br>Ala306 |
| 3 | -9.9 | Thr253<br>His413 | -7.6 | Thr252<br>Arg258<br>Glu259 | -8.6 | Thr252 | -8.1 | Thr256<br>Asn307 |
| 4 | -9.6 | Thr256 | -7.5 | Thr252<br>Thr253<br>Ala306 | -8.5 | Thr256 | -8.1 | Asn307 |
| 5 | -9.3 | No bonds | -7.5 | Trp228<br>Thr252<br>Thr256<br>Ile290 | -8.4 | No bonds | -7.9 | Thr256<br>Ala306<br>Asn307 |
| 6 | -8.8 | Thr253 | -7.1 | Thr252<br>Thr256<br>Ala306 | -8.3 | Thr252<br>Thr253 | -7.6 | Asn307 |
| 7 | -8.7 | Thr252 | -6.9 | Tyr284<br>Arg361 | -8.3 | Thr252 | -7.5 | Ala306<br>Asn307 |
| 8 | -8.4 | Arg258 | -6.8 | Glu255<br>Asn307 | -8.2 | Met192 | -7.4 | Thr256<br>Asn307 |

The best orientation of each ligand was selected for further analysis using Pymol and Ligplot+.

##### Docking of GW0742

The most stable orientation of GW0742 within the PPARβ/δ binding pocket predicted by Autodock Vina (green) was compared to the real GW0742 present in the crystal structure (pink) (**Figure 4A**). **Figure 4B** is a more detailed image showing the residues that form polar interactions with GW0742 (His247, His413 and Tyr437) according to Pymol, whereas **Figure 4C** is a 2D image created by Ligplot+ showing how the head of GW0742 forms the polar bindings and the tail is surrounded by the hydrophobic amino acids.

**Figure 4.**
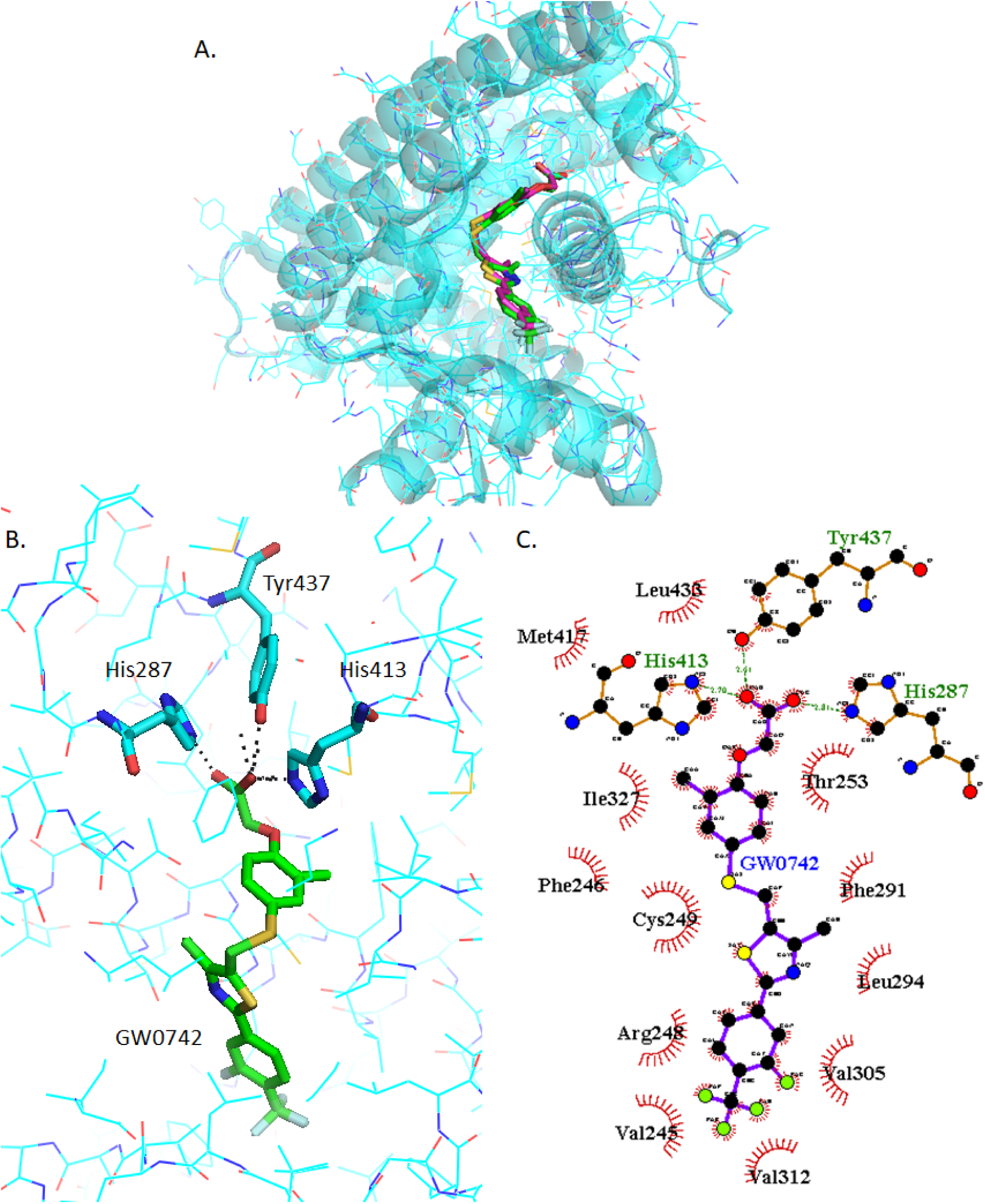
Analysis of GW0742 docked into PPARβ/δ (PBD:3TKM). A) Representation of the most stable GW0742 docking conformation (green) compared to the GW0742 of the crystal structure (pink). B) 3D detail of the amino acids forming polar bindings with GW0742 calculated by Pymol. C) Schematic 2D representation of the interaction between PPARβ/δ LBD and GW0742 calculated using Ligplot+. The green dashed lines indicate polar interactions and the red spoked arcs indicate hydrophobic interactions. Color coding of atoms: red O, blue N, mustard S, pink C of GW0742 from the crystal structure, green C of GW0742 docked into the crystal structure, cyan C from PPARβ/δ.

##### Docking of GSK3787

GSK3787 binds in a slightly different place than GW0742, although there is some overlapping of the binding sites (**Figure 5A**). Also, the amino acids involved in the polar interaction of GSK3787 predicted by Pymol, Thr252 and Asn307, are different to those of the agonists (**Figure 5B**) as well as the residues that interact with the hydrophobic tail of GSK3787 (**Figure 5C**).

**Figure 5.**
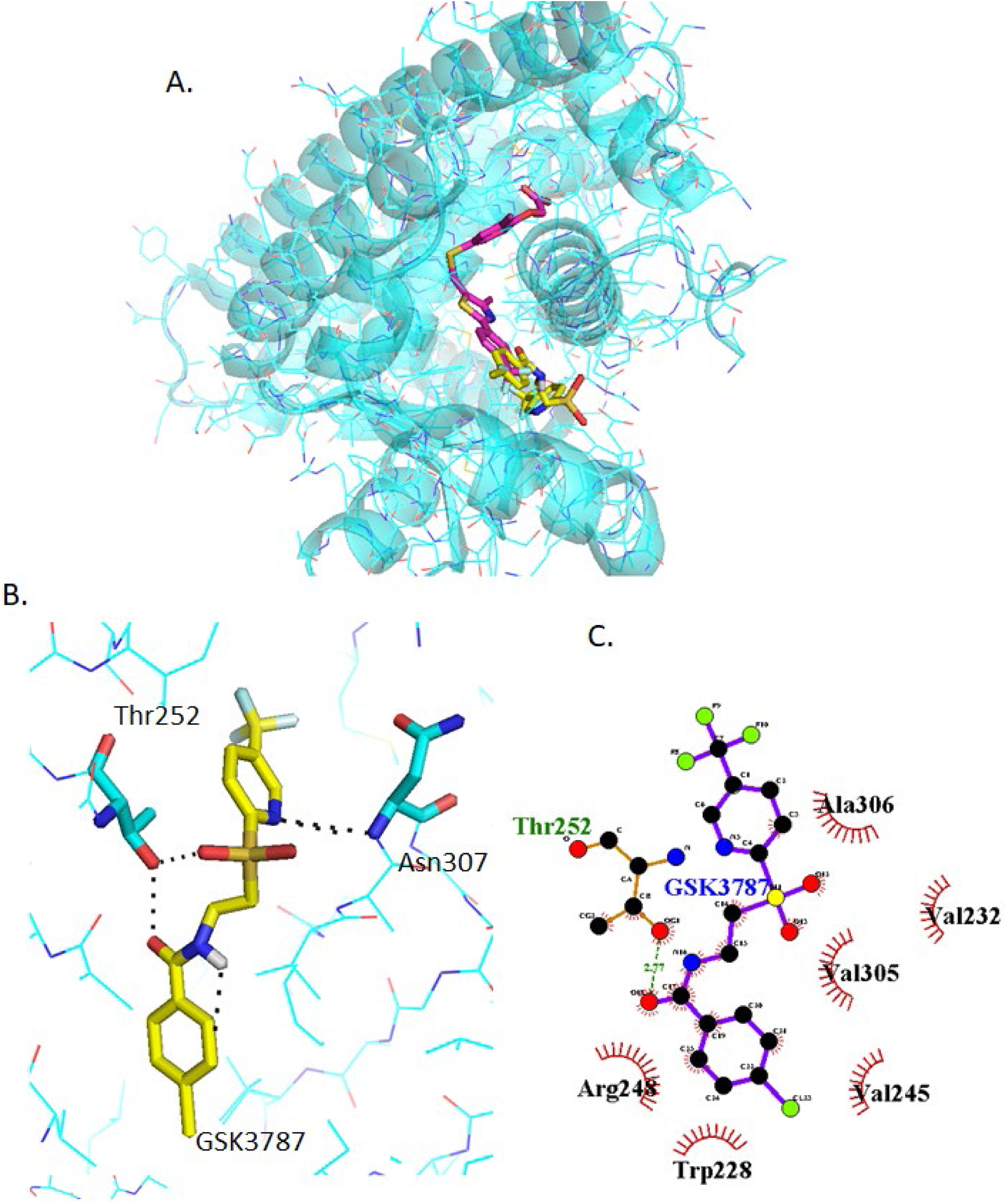
Analysis of GSK3787 docked into PPARβ/δ (PBD:3TKM). A) Representation of the most stable GSK3787 docking conformation (yellow) compared to the GW0742 of the crystal structure (pink). B) 3D detail of the amino acids forming polar bindings with GSK3787 calculated by Pymol. C) Schematic 2D representation of the interaction between PPARβ/δ LBD and GSK3787 calculated using Ligplot+. The green dashed lines indicate polar interactions and the red spoked arcs indicate hydrophobic interactions. Color coding of atoms: red O, blue N, mustard S, pink C of GW0742 from the crystal structure, yellow C of GSK3787 docked into the crystal structure, cyan C from PPARβ/δ.

##### Docking of L-165042

The most stable L-165041 orientation predicted by Autodock Vina binds in the same physical place as GW0742 (**Supplementary Figure 6A**) and the same three amino acids (His287, His413, Tyr437) form polar interactions with the head of L-165041 (**Supplementary Figure 6B**). The tail of L-165041 also forms hydrophobic interactions with a number of residues in common with GW0742, such as Val245, Arg248, Cys249, Thr253, Phe291, Leu294, Val305, Val312, Met417, Leu433 (**Supplementary Figure 6C**).

##### Docking of GSK0660

GSK0660 binds very close but not in the same binding site as GW0742 (**Supplementary Figure 7A**). The amino acids involved in the polar bindings with GSK0660, Thr252, Asn307, Arg248 and Ala306, are again different to those for the agonists, although two of them are common with GSK3787 (**Supplementary Figure 7B**). Ligplot+ predicts slightly different polar binding profile (**Supplementary Figure 7C**), probably because these two software’s use different algorithms for binding prediction, although still show hydrophobic interactions common with GSK3787, such as Trp228, Val305 and Ala306.

#### Docking of two PPARβ/δ ligands simultaneously

In order to investigate the docking of two ligands simultaneously, the first ligand was bound in the most stable orientation first. The best hit from previous docking was assigned Ligand 1, and then a second molecule was docked, assigned Ligand 2. The aim was to mimic the conditions of the experiments performed in this study and predict what could have happened at the molecular level. When the tissues were treated with only one ligand there is only one option for two ligands to bind, but when two different ligands are present at the same time either of them can bind first into the binding pocket. All these ligand-binding possibilities were considered and summarized in **Table 3**. A further analysis on Pymol was done for each option.

**Table 3.** Docking prediction of binding affinities and amino acids forming polar interactions with the PPARβ/δ ligands bound into the LBD. The affinity of the binding was predicted by Autodock Vina and the best hit was analyzed on Pymol.

| Ligand 1 |  |  | Ligand 2 |  |  |
| --- | --- | --- | --- | --- | --- |
| Molecule | Affinity (Kcal/mol) | Aa with polar interactions | Molecule | Affinity (Kcal/mol) | Aa with polar interactions |
| GW0742 | -11.1 | His287<br>His413<br>Tyr437 | GW0742 | -8.5 | Arg258 |
|  |  |  | GSK3787 | -7.7 | Trp228<br>Thr252 |
| GSK3787 | -9.1 | Thr252<br>Asn307 | GW0742 | -8.1 | Trp228<br>Lys229 |
|  |  |  | GSK3787 | -7.4 | Thr253<br>His413 |
| L-165041 | -8.7 | His287<br>His413<br>Tyr437 | L-165041 | -8.3 | Met192<br>Cys251<br>Thr252<br>Thr256<br>Ile290<br>Ala306 |
|  |  |  | GSK0660 | -6.5 | Arg198<br>Asn339 |
| GSK0660 | -8.6 | Arg248<br>Thr252<br>Ala306<br>Asn307 | L-165041 | -8.1 | Thr253<br>His413 |
|  |  |  | GSK0660 | -8.9 | Thr252<br>Thr253 |

##### Analysis of GW0742 and GSK3787 docked into GW0742-bound PPARβ/δ

Once GW0742 is bound in the most stable orientation within the PPARβ/δ-LBD, GW0742 and GSK3787 can still bind at favorable energies (−8.5 kcal/mol and −7.7 kcal/mol respectively), although at very different binding sites to the most stable one and forming polar interactions with different residues, as shown in **Figure 6**.

**Figure 6.**
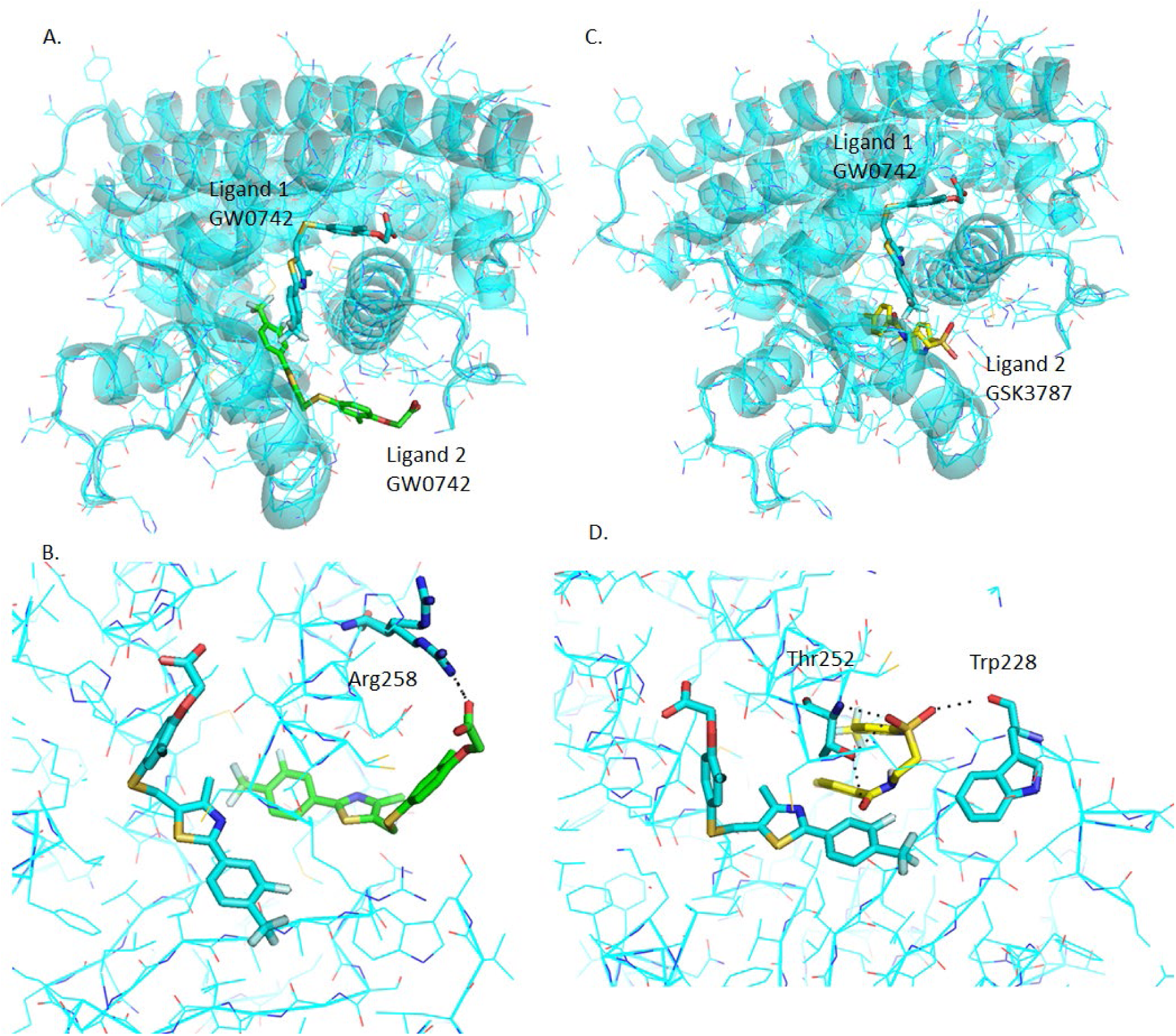
Analysis of GW0742 and GSK3787 docked into PPARβ/δ+GW0742. A second molecule of GW0742 (A, B) or GSK3787 (C, D) was docked into the LBD of PPARβ/δ containing one molecule of GW0742. A) Representation of how two GW0742 molecules bind into the PPARβ/δ binding pocket at same time. B) Detail of the amino acids interacting with the second molecule of GW0742. C) Representation of how one molecule of GW0742 first and then one molecule of GSK3787 bind into the PPARβ/δ binding pocket at same time. D) Detail of the amino acids interacting with the second molecule of GSK3787. Color coding of atoms: red O, blue N, mustard S, cyan C PPARβ/δ and GW0742 that binds first within the binding pocket, green C of GW0742 that binds second into the binding pocket, yellow C of GSK3787 that binds second into the binding pocket.

##### Analysis of GW0742 and GSK3787 docked into GSK3787-bound PPARβ/δ

GW0742 and GSK3787 can also bind into the binding pocket after GSK3787 at favorable energies (−8.1 kcal/mol and −7.4 kcal/mol respectively). The binding site is also different to the most stable ones (**Figure 7**), but interestingly the binding site is also different to the previous one, when GW0742 is bound first into the binding pocket instead of GSK3787.

**Figure 7.**
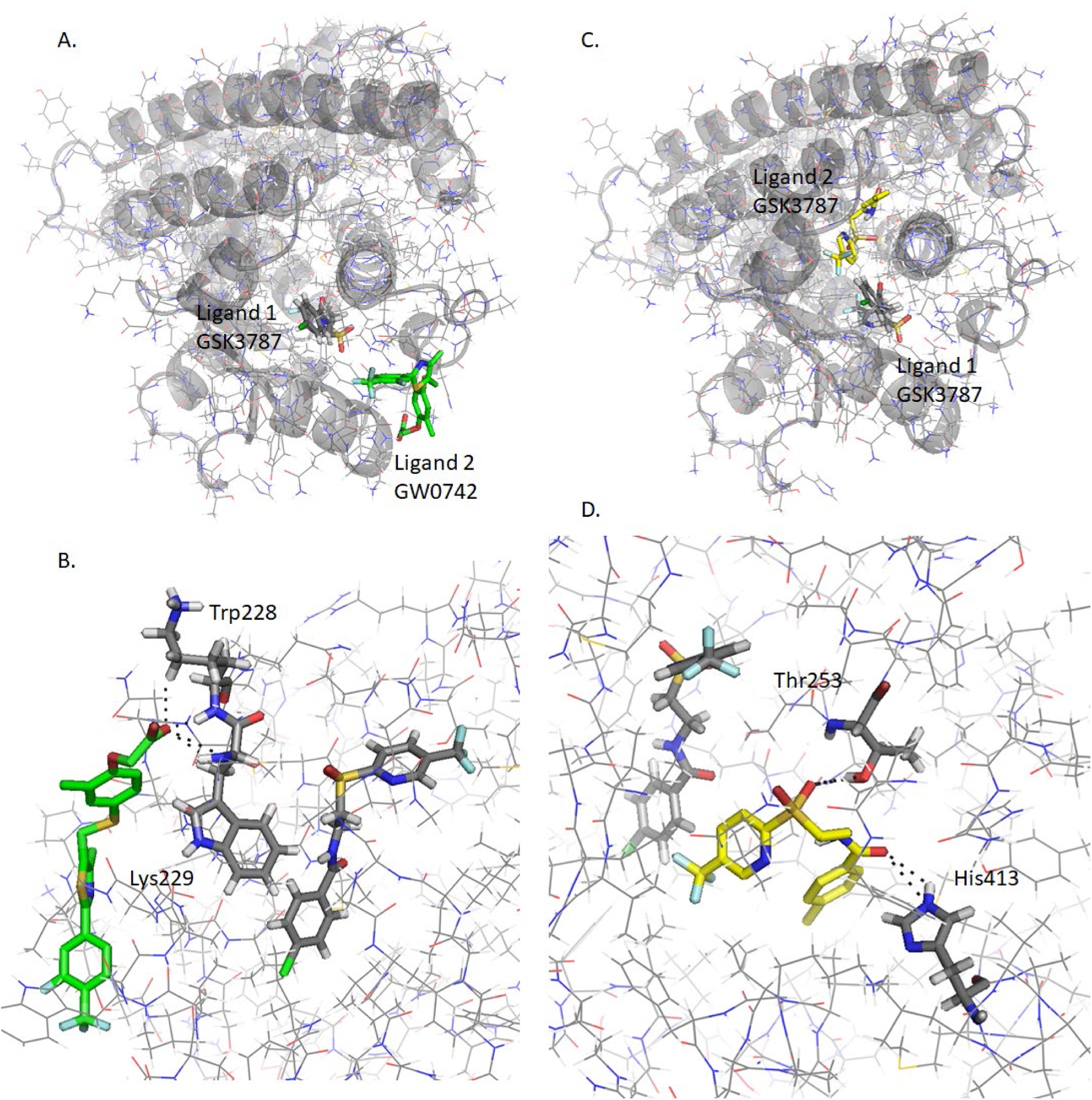
Analysis of GW0742 and GSK3787 docked into PPARβ/δ+GSK3787. A second molecule of GW0742 (A, B) or GSK3787 (C, D) was docked into the LBD of PPARβ/δ containing one molecule of GSK3787. A) Representation of how one molecule of GSK3787 first and then one molecule of GW0742 bind into the PPARβ/δ binding pocket at same time. B) Detail of the amino acids interacting with the second molecule of GW0742. C) Representation of how two molecules of GSK3787 bind into the PPARβ/δ binding pocket at same time. D) Detail of the amino acids interacting with the second molecule of GSK3787. Color coding of atoms: red O, blue N, mustard S, grey C PPARβ/δ and GSK3787 that binds first within the binding pocket, green C of GW0742 that binds second into the binding pocket, yellow C of GSK3787 that binds second into the binding pocket.

##### Analysis of L-165041 and GSK0660 docked into L-165041-bound PPARβ/δ

When L-165041 is bound to the ligand binding pocket first, a second molecule of L-165041 or GSK0660 can bind with favorable energies (−8.3 kcal/mol and −6.5 kcal/mol respectably) and again, forming polar interactions with different residues. The most interesting finding is that GSK0660, although still in the PPARβ/δ-LBD, binds outside the binding pocket (**Supplementary Figure 8**).

##### Analysis of L-165041 and GSK0660 docked into GSK0660-bound PPARβ/δ

L-165041 and GSK0660 can also bind within the binding pocket at favorable energies (−8.1 kcal/mol and −8.9 kcal/mol respectively) after GSK0660 is bound in the most stable orientation (**Supplementary Figure 9**), but again the binding site is different to previously when L-165041 was bound first.

## Discussion

This is the first report that provides insight into the PPARβ/δ molecular switch between induction and trans-repression and effects on inflammation.

PPARβ/δ is expressed in the three main tissues identified in the lung; pulmonary artery, bronchi and parenchyma, although at different concentrations. The expression of PPARβ/δ in pulmonary artery agrees with previous results where PPARβ/δ was shown to be expressed in endothelial cells (31). Bronchi express the lowest concentration of PPARβ/δ, which also agrees with the weak PPARβ/δ immunostaining of bronchial epithelial cells of mice (32), and parenchyma shows the highest expression of PPARβ/δ, which differs to the expression profile in mice, where the expression of PPARβ/δ is also weak in alveolar type II cells (32).

The effects of PPARβ/δ ligands on LPS-induced inflammation of the tissues was studied by incubating the tissues with different agonists (GW0742 and L-165041) and antagonists (GSK3787 and GSK0660), which showed similar regulatory profile in all tissues. In general terms, LPS induces the release of NO and IL-6, and the incubation with one ligand (either agonist or antagonist) does not reduce the production of these inflammation markers. However, co-incubation with both an agonist and an antagonist significantly reduces LPS-induced inflammation.

However, it cannot be ignored the fact that these results disagree with a large number of studies where the treatment with GW0742 attenuated the lung inflammation in murine models, the effects of GW0742 were abolished or significantly reduced by GSK0660, and the PPARβ/δ deletion exacerbates the lung inflammation (33-35). The difference is that these studies are *in vivo* and, although the inflammation is monitored in the lung, the whole organism is responding to the infection, the immune system is being activated and immune cells such as neutrophils and macrophages are infiltrated into the alveolar cavity. This means that there is a synergic response composed of different tissues and cells that makes the anti-inflammatory response more effective. In the model used here, the tissues were separated from each other and the LPS-induced regulation of inflammation by PPARβ/δ was analyzed in isolated tissues to investigate our molecular switch hypothesis. Under these conditions, PPARβ/δ seems to play a more important role in the regulation of inflammation in the pulmonary artery than in bronchi and lung parenchyma.

The initial hypothesis was that PPARβ/δ acts as a molecular switch between induction and trans-repression and depending on which mechanism is triggered it will have pro- or anti-inflammatory effects. If the hypothesis is true, the results would indicate that the incubation with an agonist led to a pro-inflammatory state of PPARβ/δ and the incubation with the agonist and antagonist together switches PPARβ/δ to an anti-inflammatory state. However, the data do not provide enough information to conclude whether the pro- and anti-inflammatory response is regulated through the induction or trans-repression mode of action. With this aim, qRT-PCR was performed for the expression of marker genes for each mechanism of action in all treatments and all tissues.

Marker genes were selected to represent the induction and trans-repression modes of action. Khozoie (36) and Adhikary (37) performed a genomewide analysis of genes regulated by PPARβ/δ in mouse keratinocytes and human myofibroblasts respectively. Khozoie (36) cross-linked the two lists of genes regulated by PPARβ/δ and created a new list of 103 genes regulated by PPARβ/δ in both human and mouse models. There is a high possibility that these genes are regulated by PPARβ/δ in rats as well, therefore this list was used to select the induction and trans-repression marker genes.

It is very tempting to accept that those genes found with PPARβ/δ bound to its promoter are regulated through the induction mode of action and those without PPARβ/δ bound to the promoter are regulated through the trans-repression mode of action, however our results indicate that this is incorrect. For example, *Angptl-4* is well known to be regulated by induction mode (38-40), which agrees with Khozoie (36) results, where they found PPARβ/δ bound to the promoter of *Angptl-4*. However, *Pdk-4* is another well-known gene regulated in the induction mode (39-41), but Khozoie (36) did not identify PPARβ/δ bound to the promoter of the gene. Consequently, it can be presumed that the genes with PPARβ/δ bound to the promoter are regulated through the induction mode of action, but it cannot be ruled out that genes without PPARβ/δ in their promoter are regulated in the trans-repression mode. Therefore, other ways of identifying potential markers for the trans-repression mode of action are needed.

A separate study was used to identify the genes regulated by NF-κB after LPS activation of human cells (42) and crosslinked with the list of 103 genes regulated by PPARβ/δ in mouse and human. After LPS-activation, NF-κB was found in the promoter of four genes common in the two lists, whose transcription was also increased (42). Two of them were selected as potential markers for the trans-repression mode of action of PPARβ/δ: *Timp-1* and *Serpine-1*. The idea is that when PPARβ/δ is activated and functioning in the trans-repression mode of action it will sequester NF-κB and inhibit the transcription of the selected genes.

Bcl-6 has shown to be another important trans-repression mechanism of action of PPARβ/δ, for that reason another marker gene for this pathway was identified. One study showed that LPS increases the transcription of *Id2* in some tissues, like the brain (43), and another study showed that *Id2* is regulated by Bcl-6 (44), therefore *Id2* was chosen as a marker gene of the PPARβ/δ trans-repression mode of action through Bcl-6. When PPARβ/δ is not activated it sequesters Bcl-6, which will allow the transcription of *Id2* in the presence of LPS; should PPARβ/δ become activated it will release Bcl-6 which will then repress the expression of *Id2*. Our study indicates that Bcl-6 response was not prominent in the LPS response in any lung tissue studied.

The qRT-PCR of pulmonary artery showed that the activation of PPARβ/δ with GW0742 increases the transcription of *Pdk-4* and *Angptl-4* and represses the transcription of *Timp-1* and *Serpine-1*, indicating that agonist activation of PPARβ/δ triggers both, induction and trans-repression. However, the presence of GW0742 and GSK3787 together does not induce the transcription of *Pdk-4* or *Angptl-4* but it still represses the transcription of *Timp-1* and *Serpine-1*, indicating that the trans-repression but not induction is occurring. The transcription of *Id2* seems not to be affected by any of the treatments used in this study and therefore it can be concluded that *Id2* is not regulated by PPARβ/δ in pulmonary artery.

By combining these findings with the inflammatory response, it can be deduced that in pulmonary artery GW0742 triggers both induction and trans-repression, resulting in a pro-inflammatory state of PPARβ/δ. Interestingly, the presence of GW0742 and GSK3787 together inhibits the induction mode of action but activates the trans-repression mode of action through NF-κB, which is the treatment that produces the strongest anti-inflammatory effect. Taken together, it suggests that GW0742 triggers the induction mechanism, which is pro-inflammatory, and GW0742 plus GSK3787 switches PPARβ/δ to the trans-repression mechanism through NF-κB which is responsible for the anti-inflammatory effects of PPARβ/δ in LPS-induced inflamed pulmonary artery.

In bronchi, *Angptl-4* is not expressed but *Pdk-4* is significantly increased by GW0742. On the contrary, in lung parenchyma *Angptl-4* is significantly increased by GW0742 but *Pdk-4* is not affected by PPARβ/δ ligands. This result suggests that the induction mechanism of action of PPARβ/δ is occurring in bronchi and lung parenchyma. On the other side, the transcription of *Timp-1, Serpine-1* and *Id2* does not vary with the treatments, indicating that trans-repression mode is not a major mechanism of action of PPARβ/δ in these tissues, which can explain the weaker inhibition of the inflammatory response.

It has been shown in several studies that *Nos2* is regulated by PPARβ/δ (35,45,46), although it has not been described whether this regulation is via induction or trans-repression. In our hands, LPS increases *Nos2* expression in all tissues tested, however the regulation of its expression by PPARβ/δ is different depending on the tissue. In pulmonary artery, GW0742 represses the expression of *Nos2*, suggesting that PPARβ/δ is inhibiting *Nos2* through the induction mode of action. Surprisingly, in bronchi GW0742 significantly increases the transcription of *Nos2* compared to LPS, suggesting that PPARβ/δ enhances the expression of *Nos2* through the induction mode of action, and in parenchyma the expression of *Nos2* is not regulated by PPARβ/δ. This result indicates that the cell/tissue type is another important co-determinant of the type of response of the PPARβ/δ target genes – different co-regulators might be available depending on the tissue, resulting in alteration of the dynamics of transcriptional complexes and interactions with DNA binding sites. Thus, there are multiple levels of regulation by which PPARβ/δ can influence the expression of target genes.

Further to this, It has been suggested that the large ligand binding pocket of PPARβ/δ can accommodate more than one ligand, resulting in an unusual PPAR:ligand stoichiometries that could trigger inactive receptor conformations (28), a possibility that was further investigated using *in silico* methods.

Firstly, the two PPARβ/δ agonists (GW0742 and L-165042) and two antagonists (GSK3787 and GSK0660) were docked into the PPARβ/δ binding site. It was found that the agonists and antagonists have a different binding profile within the binding pocket: the same three amino acids His287, His413 and Tyr437 form polar interactions with the two agonists tested – but they do not bind the antagonists – and the amino acids Thr252 and Asn307 are more prone to bind the antagonists.

This finding agrees with previous results were GW0742 was docked to another PPARβ/δ crystal structure (PDB: 3GZ9) using another docking software (Glide), and the same three amino acids bound to GW0742 (47). Furthermore, several studies co-crystallized PPARβ/δ with different agonists both synthetic, such as iloprost (48), the fibrate GW2433 (49), or GW501516 (50) and natural PPARβ/δ agonists such as with eicosapentaenoic acid (EPA) (49), and in all cases the agonists showed polar bindings with the same three amino acids His287, His413 and Tyr437. It is worth mentioning another study where the authors selected 5 compounds that potentially bound PPARβ/δ and performed a luciferase transactivation assay to biologically test if these compounds activate PPARβ/δ. They further analyzed two of them by docking and molecular dynamics (MD) simulation, one compound that activated PPARβ/δ (Compound 1) and another one that did not activate PPARβ/δ (Compound 2). The docking and MD simulation results for the Compound 1 showed an interaction with His287, His413 and Tyr437, and the results for Compound 2 showed an interaction with Thr252 (51).

This suggests the possibility that the different binding profile between agonists and antagonists can provoke a different 3D conformational change that might explain why PPARβ/δ binds to co-repressors instead of co-activators and *vice versa*. Taking all this into account it can be hypothesized that a ligand that shows a high binding affinity and is predicted to form polar bonds with His287, His413 and Tyr437 will most likely behave as agonist. On the contrary, if one ligand shows high binding affinity but it is predicted to bind other residues such as Thr252 and Asn307 it is more likely that it will behave as antagonist.

Next, the possibility of two ligands binding into the PPARβ/δ binding pocket at same time was investigated. The treatment of tissues with only GW0742 allows two possibilities: the binding of one or two molecules into LBD. If a second molecule of GW0742 binds to PPARβ/δ, this molecule is predicted to bind not too far from the most stable binding site and with the same binding affinity and residue interaction (Arg258) than the 8^th^ best pose predicted for the first molecule. Similarly, when only GSK3787 is present in the treatment, a second molecule of GSK3787 is predicted to bind also not far away from its most stable binding site and with favorable binding affinity, and what is more, it will still form polar bonds with Thr252, an amino acid that is predicted to bind GSK3787 in five out of eight most stable poses predicted by docking, including the first three.

For the treatment of tissues with GW0742+GSK3787 all the options mentioned above still apply but two more options are available: GW0742 binds first and GSK3787 after or GSK3787 binds first and GW0742 after. When GW0742 binds first, GSK3787 can still bind very close to its most stable binding site with a very favorable binding affinity, and what is more, still binds the residue Thr252. If GSK3787 binds first, GW0742 is predicted to bind very far away from the most stable binding site, at the entrance of the binding pocket, and as a consequence it will have a very different binding profile forming polar bonds with Trp228 and Lys229, two residues that did not show any interaction with ligands before. Similar analysis can be done with the other pair of agonist and antagonist L-165041 and GSK0660.

Taking into account the docking scores and molecular poses of the ligands, all possibilities described above have very favorable energies for it to happen. That opens a whole new scenario of possibilities that could dramatically change the 3D conformation of PPARβ/δ in ways that have not been thought of before, resulting in the binding of different co-regulators, which ultimately could change the PPARβ/δ response from induction to trans-repression or *vice versa*. To our knowledge, this is the first time that the possibility of binding two ligands simultaneously into the PPARβ/δ binding pocket has been explored. The results suggest that this possibility is very likely to happen with very favorable affinity energies, and it is worth considering when designing and interpreting experiments where PPARβ/δ is ligand-activated.

In summary, this is a multidisciplinary approach of the study of PPARβ/δ that provides novel information about its functioning at molecular level. In the model used here the presence of agonist and antagonist at same time switches the PPARβ/δ mode of action from induction to trans-repression, and the trans-repression mode of action was linked with anti-inflammatory effects. Among the tissues tested, PPARβ/δ seems to play a more important role in the regulation of inflammation in the pulmonary artery. A characteristic PPARβ/δ-ligand binding profile which is different for agonists and antagonists is also described. PPARβ/δ agonists form polar bonds with His287, His413 and Tyr437, while antagonists are more promiscuous about which amino acids they bind to, although they are very prone to bind Thr252 and Asn307. The possibility of two agonists binding at same time into the PPARβ/δ binding pocket was also explored, and all the options studied seem feasible with favorable binding energies, suggesting the need for caution when designing and interpreting the results of experiments using PPARβ/δ ligands.

## Experimental procedures

### Reagents

The PPARβ/δ ligands GW0742, GSK3787, L-165042 and GSK0660 as well as 1400W, LPS O55:B5, sulfanilamide and naphthylethylenediamine dihydrochloride were purchased from Sigma. Sodium nitrate, orthophosphoric acid and DMSO were purchased from Fisher Scientific. Primers from Applied Biosystem: β-actin (Rn00667869_m1), Pdk-4 (Rn00585577_m1), Angptl-4 (Rn015228817_m1), Nos2 (Rn00561646_m1), Serpine-1 (Rn01481341_m1), Timp-1 (Rn01430873_m1), Id2 (Rn01495280_m1).

### Animals

Male Wistar rats (350-450 g) were sourced from Charles River (UK) and housed in pairs in standard cages (Tecniplast 2000P) with sawdust (Datesand grade 7 substrate) and shredded paper wool bedding with water and food (5LF2 10% protein LabDiet) in the Biological Services Unit at the University of Hertfordshire. The housing environment was maintained at a constant temperature of 22 ± 2 °C, under a 12 h light/dark cycle (lights on: 07:00 to 19:00 h). All testing was conducted under the light phase of the animals’ light/dark cycle, and care was taken to randomize treatment sequences to control for possible order effects.

All experiments were approved by the local ethics committee, and conducted in accordance with the guidelines established by the Animals (Scientific Procedures) Act, 1986 and European directive 2010/63/EU. Rats were euthanized according to schedule 1 procedure by CO_2_ asphyxiation followed by cervical dislocation. Lungs were removed and immediately placed in physiological saline solution (PSS) buffer (118 mM NaCl, 4.7 mM KCl, 2.5 mM CaCl_2_, 1.17 mM MgSO_4_, 1 mM KH_2_PO_4_, 5.5 mM, glucose, 25 mM NaHCO_3_ and 0.03 mM Na_2_EDTA).

Following dissection, tissues were incubated in 1% v/v penicillin/streptomycin in DMEM under 5% v/v CO_2_ at 37 °C in the required treatment (detailed in **Table 1**) for up to 24 hours. After incubation, the culture medium was removed and stored at −20 °C until Greiss assay or IL-6 ELISA analysis. The tissues were stored at −80 °C until needed for mRNA extraction for qRT-PCR.

### Quantification of PPARβ/δ expression in lung tissues by enzyme-linked immunosorbent assay (ELISA)

The expression of PPARβ/δ on pulmonary artery, bronchi and parenchyma from naïve rats was measured using Rat PPARβ/δ ELISA kit (Abbkine) according to the manufacturer’s instructions. Briefly, the dissected tissues were homogenized with liquid N_2_ using a mortar and a pestle, and the proteins were extracted in ice-cold phosphate buffered saline (PBS) with proteinase inhibitor cocktail in a ratio 9 mL PBS (137 mM NaCl, 2.7 mM KCl, 10 mM Na_2_HPO_4_, 1.8 mM KH_2_PO_4_, pH 7.4) per g tissue. The samples were then sonicated 3×30 seconds and centrifuged for 5 min at 5000 g. The supernatant was collected, and the protein concentration was quantified by BCA assay. Then, 50 µL of standards and samples were added in duplicate to the plate containing pre-coated anti-PPARβ/δ and incubated 45 min at 37 °C. The wells were washed and incubated with 50 µL of the horseradish peroxidase (HRP)-conjugated detection antibody for another 30 min at 37 °C. The wells were washed again and incubated in the dark with the chromogen solution for 15 min at 37 °C. Finally, the reaction was stopped by adding 50 µL of Stop solution and the plate was immediately measured in a microplate reader. The concentrations of PPARβ/δ were determined by comparison of OD_450_ to a standard curve (0-8 ng/mL).

### Quantification of nitric oxide released by lung tissues by the Griess assay

An aliquot of the culture medium (50 µL) thawed on ice was mixed with an equal volume of Griess reagent (mixture of equal volumes Griess reagent 1 and Griess reagent 2 containing sulfanilamide 1% w/v + orthophosphoric acid 5% v/v and naphthylethylenediamine dihydrochloride 0.5% w/v respectively). The concentration was determined by comparison of the OD_540_ to a standard curve of solutions of sodium nitrite (0-1 mM).

### Quantification of IL-6 released by lung tissues by ELISA

The release of IL-6 by lung tissues to the culture medium was measured using Rat IL-6 DuoSet ELISA kit (R&D Systems) according to the manufacturer’s instructions. Briefly, all samples and standards were conducted in duplicate to a microtiter plate containing the capture antibody. After two hours of incubation the wells were washed and incubated with the detection antibody for another two hours. The wells were washed again and incubated in the dark with Streptavidin-HRP for 20 min. The wells were washed once more and incubated in the dark with Substrate Solution. After 20 min the stop solution was added and immediately measured in a microplate reader. The readings at 540 nm were subtracted from the readings at 450 nm and the concentrations of samples were determined by comparison with the standard curve (0-8 ng/mL).

### Quantitative real time-polymerase chain reaction (qRT-PCR)

Total RNA was extracted from tissues using RNeasy Fibrous Tissue Mini Kit (Quiagen). The tissues were first pulverized in a pestle with liquid N_2_ and the RNA was then extracted following the manufacturer’s instruction and stored at −80 °C until use. The quality and concentration of the RNA was measured using Nanodrop (SimpliNano, GE Healthcare Life Science) at a wavelength of 260 nm. Further to that, the degradation of RNA was checked in a 1% w/v agarose gel, and the DNA contamination of the RNA was checked by PCR. To do that, genomic DNA was also extracted from the tissues using PureLink Genomic DNA mini kit from Invitrogen following the manufacturer’s instructions and used as a positive control of the PCR, and two primers for the housekeeping gene *β-actin* were designed (forward CTGGTCGTACCACTGGCATT, reverse AATGCCTGGGTACATGGTGG). The termociclator Mastercycler Nexus gradient (Eppendorf) was set with the following PCR protocol: 95 °C for 10 min, 40 cycles of 95 °C for 30 s, 56 °C for 30 s and 72 °C for 30 s, 72 °C for 10 min and hold at 4°C.

cDNA was obtained by reverse transcription (RT) using iScript cDNA synthesis kit (Bio-Rad) following the manufacturer’s instructions. The RT was performed using a thermociclator Stratagene Mx3005P (Agilent Technologies) with the following steps: 5 min at 25 °C, 20 min at 46 °C, 5 min at 95 °C, and hold at 4 °C.

qRT-PCR was performed to analyze mRNA expression using a Taqman System. Briefly, 10 µL of reaction mix containing the primers and cDNA was incubated in a 96 well-plate following the cycle conditions: 95 °C for 10 min, 40 cycles of 95 °C for 15 s and 60 °C for 1 min.

### Docking

The ability of drugs to bind into protein active sites was investigated using AutoDock/Vina with Pymol and Ligplot+ as a graphical user interface. For the docking simulations, the PPARβ/δ crystal structure 3TKM was selected for having one of the highest resolutions (1.95 Å). The PDB file was downloaded from Protein Data Bank. Water molecules, ligands and other hetero atoms were removed from the protein structure, and the addition of hydrogen atoms to the protein was performed using AutoDock Tools version 1.5.6. The grid was set manually to cover the active site. The file was saved as a pdbqt file.

The ligand molecule structures were drawn in ChemSketch, the energy was minimized and saved in PDB format, and converted into a pdbqt file with AutoDock Tools version 1.5.6. Molecular docking was performed with the software AutoDock Vina and all parameters set as default. Results with minor calculated free energy variations were analyzed using Pymol version 1.7.4 and LigPlot+ version1.4.5 softwares.

For the docking of two molecules, the 3TKM PDB file without hetero atoms was combined with the best docking result of each ligand in one single PDB file, one PDB file per ligand. These files were opened in Autodock Tools, H_2_ were added, the grid was set manually and saved in a new pdbqt file. This file was used for the docking with the second molecule.

### Statistical analysis

Statistical comparisons were performed on GraphPad Prism 5.0 software using one-way or two-way ANOVA with Bonferroni’s post hoc analysis for NO and IL-6 detection assays, and Dunnett’s post hoc analysis for qRT-PCR. The values are expressed as observed mean ± SEM. Data was normalized previously to the statistical analysis. In short, data from NO and IL-6 detection assays was normalized against the group treatment LPS and expressed as a fold change. The relative quantification of genes analyzed by qRT-PCR was calculated with the comparative CtΔΔ method, β-actin was used as endogenous control and data was normalized against the control group.

Values of p<0.05 were considered statistically significant. When the level of probability (P) is less than 0.05 (*), less than 0.01 (**) or less than 0.001 (***), the effect of the difference was regarded as significance.

